# Shared striatal neurons exhibit context-specific dynamics for internally and externally driven actions

**DOI:** 10.1101/2025.09.18.677137

**Authors:** Jan L. Klee, Sulekh Fernando-Peiris, Sahil Suresh, Ines Rodrigues-Vaz, Darcy S. Peterka, Rui M. Costa, Vivek R. Athalye, Tanya Sippy

**Author notes:** These authors contributed equally to the work (Jan L. Klee, Sulekh Fernando-Peiris).

## Abstract

Animals can initiate movements either in response to external cues or from internal drive, yet how the brain flexibly supports both remains unclear. Disorders such as Parkinson’s disease disrupt these modes differently, suggesting distinct underlying mechanisms. Such differences could arise from specialized circuits dedicated to each action mode or from shared neuronal populations that shift their dynamics across contexts. To distinguish between these possibilities, we performed two-photon calcium imaging in the dorsolateral striatum as mice executed the same lever press either spontaneously or in response to a cue, enabling direct comparison of internally and externally triggered actions. Unsupervised clustering revealed subpopulations of neurons modulated during cue, movement, or post-action periods. Critically, the same neurons were tuned to movement across initiation contexts. However, the dynamics of this shared population evolved within distinct “context” and “action” subspaces, even before movement initiation. Both dopamine D1- and D2-receptor-expressing spiny projection neurons contributed to both subspaces, with D1-SPNs more active at the time of the sensory stimulus. These results show that context shapes neural dynamics within a shared movement-encoding population, revealing a context-generalizable striatal code that supports flexible movement initiation across internal and external drives.

## INTRODUCTION

Understanding the mechanisms that distinguish self-initiated from sensory-evoked movements is essential for elucidating the neural basis of motor control. This distinction has clinical relevance: patients with Parkinson’s disease often experience difficulty initiating voluntary movements, yet can transiently regain motor function in response to external cues such as visual or auditory stimuli (*1*, *2*). These observations suggest that sensory-guided movements engage neural circuits that are at least partially distinct from those underlying internally generated actions. At the cortical level, self-evoked and externally triggered actions are represented by largely distinct neural ensembles (*3–5*) —for example, sensory cortical neurons are preferentially recruited by external cues, whereas medial frontal neurons show ramping activity before self-initiated movements. However, recent population-level analyses point to a shared preparatory component within these areas, indicating that overlapping neuronal populations can support both initiation modes (*6*). More broadly, prior work has demonstrated that cortical circuits often embed multiple task variables within shared low-dimensional subspaces (*7*, *8*), suggesting that mixed selectivity and population-level organization may allow flexible routing of both self-initiated and stimulus-driven signals.

The striatum, the input nucleus of the basal ganglia, integrates information from widespread cortical and subcortical sources (*9–13*), enabling context-dependent initiation of the same action (*14*, *15*). Its cortical and subcortical inputs are topographically organized: anterior regions of the dorsal striatum are predominantly innervated by motor areas, whereas more posterior regions receive dense input from sensory areas, including visual, somatosensory, and auditory cortex (*16–19*). In the dorsolateral striatum (DLS) individual neurons are thought to receive convergent input from both sensory and motor cortical areas (*20–22*), positioning them to integrate contextual and motor-related signals. Consistent with this, single-unit recordings in non-human primates demonstrated that subsets of striatal neurons are preferentially engaged during either self-initiated or stimulus-driven movements, while only a minority show overlapping activity across both contexts (*23*).

Striatal projection neurons (SPNs) are broadly divided into two major classes based on their dopamine receptor expression: D1-SPNs, which form the direct pathway and are classically associated with promoting action initiation, and D2-SPNs, which form the indirect pathway and are linked to action suppression (*24*). While these pathways have traditionally been viewed as functionally opposing, accumulating evidence indicates that both are active during movement and may contribute in complementary ways to shaping behavior (*25–28*). Recent work from our group and others has shown that in sensory-driven behaviors, D1-SPNs often exhibit stronger and more time-locked activation to external cues (*29–33*), suggesting that the direct pathway may be particularly responsive to salient sensory events that trigger action.

However, it remains unclear how these inputs are funneled to drive action in the appropriate context, and how D1- and D2-SPNs may differentially contribute to this process. Do distinct contextual demands recruit non-overlapping neuronal populations, or do shared neurons integrate different inputs depending on the mode of initiation? Understanding how context-specific inputs are transformed into action within striatal circuits is essential for revealing the mechanisms of flexible motor control. At one end of the spectrum, neurons in the dorsal striatum may form entirely distinct populations that separately encode cues and actions, while at the other, the same neurons may exhibit overlapping activity that integrates both types of information, potentially along different population activity dimensions.

To address this question, we designed a task in which mice performed a simple, well-defined movement either spontaneously or in response to an external cue, enabling direct comparison of internally generated and stimulus-driven actions. By holding motor output constant, we isolated striatal activity differences that reflect initiation mode rather than movement kinematics, revealing how the striatum contributes to self-initiated versus cue-guided behavior. Two-photon calcium imaging in the dorsolateral striatum (DLS) showed that, through unsupervised hierarchical clustering, neurons segregated into subpopulations tuned to the cue, movement, or post-movement epochs. One cluster was consistently active around movement regardless of initiation mode, identifying a shared action-timed ensemble. Despite this overlap, population dynamics diverged before movement onset, and trial type could be reliably decoded using support vector machines. Within the same population, subspace analyses revealed orthogonal components encoding initiation context and movement execution. Both D1- and D2-SPNs participated in these subspaces—D1-SPNs showing elevated activation near cue onset and movement initiation, and D2-SPNs exhibiting increased post-action activity across both trial types. These findings show that internal and external drivers of behavior converge onto shared action-encoding neurons, whose context-dependent dynamics define a flexible population code for action within basal ganglia circuits.

## RESULTS

### Alternating cue-evoked and self-paced lever task

We first developed a task in which mice learn to perform a self-paced and cue-evoked action within the same behavioral session, in different blocks. Food deprived mice first learned to push a lever to receive a sucrose reward (Fig. S1A). On subsequent training days, they learned to push the lever in response to an auditory cue (5 kHz, 100 ms pure tone). On the day of behavioral testing, mice performed these two behaviors in three sets of blocks, each of which began with a “cued block” followed by “self-paced block” (Fig. 1A-B), with block transitions triggered after ∼30 rewards were obtained. In both block types, the rewarded lever trajectories were highly similar (Fig. 1C-E). In cued blocks, mice pushed the lever in a manner that was time locked with the cue while in self-paced blocks, lever pushes occurred at variable times after the trial start which resulted in significantly shorter “reaction times” during the cued blocks (Fig. 1F, reaction time: cued = .30 s, self-paced = 6.0 s; Wilcoxin signed rank test, p = 0.03, n = 6). The rate of lever pushes was similar across block number within a block type (Fig. 1G, cued = 16.9 pushes/minute; self-paced = 16.0 pushes/minute; Wilcoxin signed rank test, p = 0.69, n = 6), as was the rate of rewarded lever pushes (Fig. 1H, cued = 7.49 rewards/minute; self-paced = 7.0 rewards/minute; Wilcoxin signed rank test, p = 0.84, n = 6). Finally, the ratio of rewarded lever presses to all presses was not significantly different across the trial blocks in either block type (Fig. 1I, cued = 0.50; self-paced = 0.45 ; Wilcoxin signed rank test, p = 0.68, n = 6). This consistency in behavior across blocks indicates that performance was stable over time, allowing us to attribute differences between block types to task structure rather than changes in motivation or motor output. Importantly, inhibition of neural activity in the dorsolateral striatum with fluorescent muscimol significantly decreased task performance during both self-initiated behavioral sessions and cue-evoked sessions (Fig. S1F-H). This was in contrast to inhibition of the posterior, or “tail” of the striatum in which fluorescent muscimol specifically had an effect on cue-evoked lever pressing (Fig. S1I-K). This finding highlights a functional dissociation along the anterior-posterior axis of the striatum, suggesting that the dorsolateral striatum is broadly necessary for task performance regardless of initiation context, whereas the posterior striatum plays a more selective role in mediating cue-driven actions.

**Figure 1:**
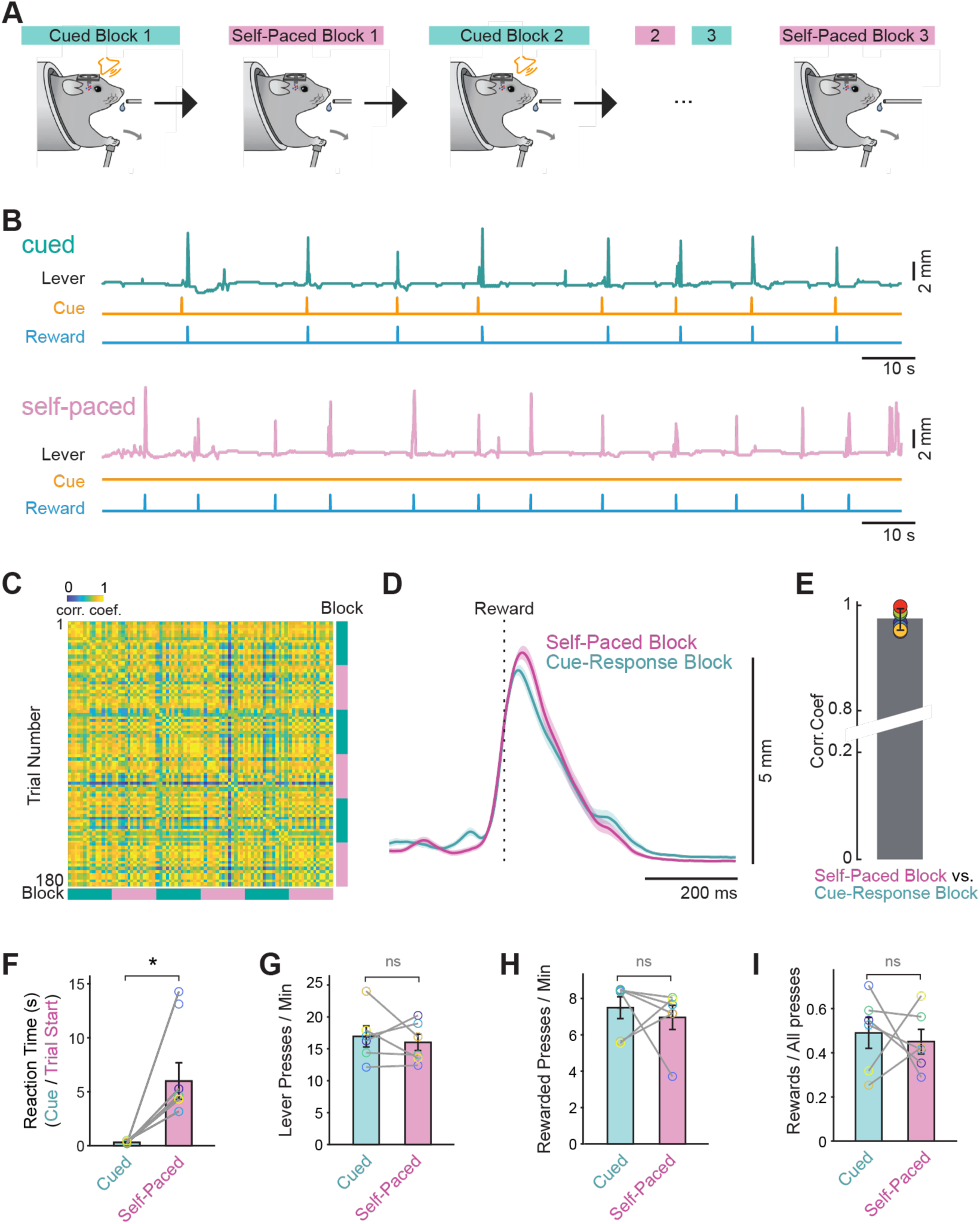
The “switching” task allows for the assessment of a self-paced and cue driven actions within a single session. (A) Schematic of the switching task depicting alternating block design within the same behavioral session (B) Top: example of lever trace in a cued block. Bottom: example of a trace in a self-paced block from this same mouse. (C) Cross correlation matrix of all rewarded lever push trajectories from an example mouse across all blocks during the test day. (D) Average lever press trajectories for self-paced and cued lever push trajectories for the same mouse as in C. (E) The correlation coefficient between self-paced and cues lever push trajectories for all mice, plotted at mean ± sem. (F) The reaction time after cue onset (cued blocks) and trial onset (self-paced blocks) was statistically significantly different in the two clock types (Wilcoxin signed rank test: * p < 0.05). (G) The lever push frequency (per min) was not different between cued and self-paced blocks (Wilcoxin signed rank test: ns > 0.05). (H) The rate of rewarded pushes (per min) was not different between cued and self-paced blocks (Wilcoxin signed rank test: ns > 0.05). (I) The fraction of lever pushes that led to reward was not different between cues and self-paced blocks (Wilcoxin signed rank test: ns p > 0.05). All bar plot data are represented as mean ± SEM. Each open circle represents an individual mouse.

### A shared movement-timed DLS ensemble emerged across contexts

To investigate the neural activity underlying cue-evoked and self-paced lever pushing, we virally expressed the Ca^2+^ indicator GCaMP8m (n = 4) or GCaMP8f(*34*) (n =2) in the dorsolateral striatum (DLS, Fig. 2A). To distinguish between D1- and D2-SPNs, we performed experiments in mice genetically engineered to express tdTomato in D2-SPNs or D1-SPNs (generated by crossing D2-Cre or D1-Cre mice(*35*, *36*) with Ai14reporter mice(*37*)) (Fig. 2B-C). Imaging commenced on the first day they were required to alternate between cue-evoked and self-paced lever blocks, beginning with a cue block in all mice, as described above. Individual ROIs were semiautomatically extracted using Suite 2P(*38*) and showed behaviorally relevant fluorescence changes (Fig. 2D-E). We then took the grand trial average z-scored deltaF/F responses in self-paced and cue-response blocks and aligned them to the time of the lever push (Fig. 2F, dark teal and magenta traces) and found them to be similar in the 1 s leading up to the push action. Notably, when we aligned the trials in the cue-evoked blocks to the cue, we observed a sharp rise in average activity that coincided with the onset of the cue (Fig. 2F, light teal trace).

**Figure 2:**
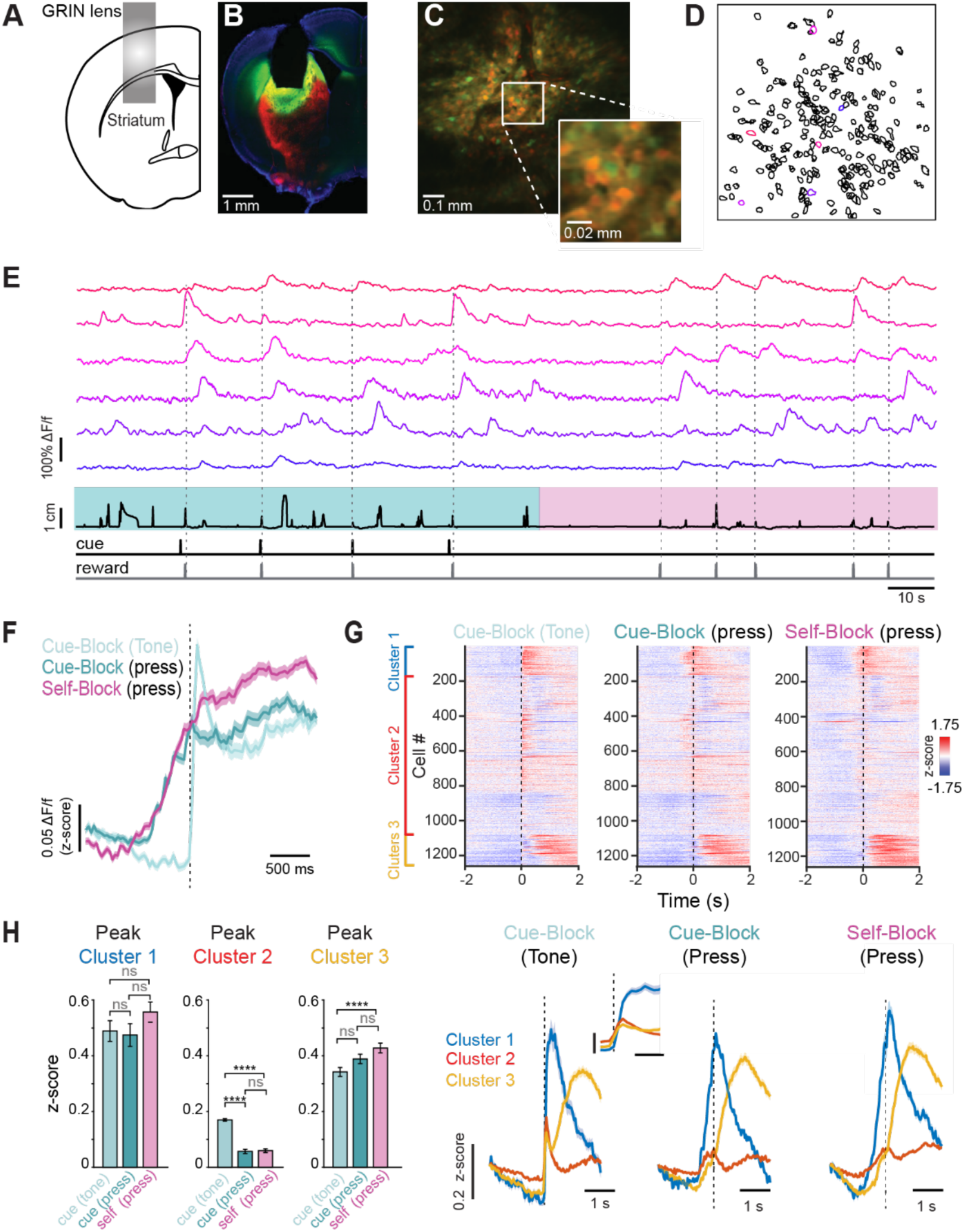
Population imaging of DLS shows common motor representations alongside context-dependent ensembles. (A) Schematic of GRIN lens position in the DLS. (B) Example of histological verification of GRIN lens placement and GCaMP8m expression in a D2-tdTomato mouse (C) Field of view (FOV) through the GRIN lens in one example mouse (D) Regions of interest (ROIs) identified from one switching task session of the same mouse in C. Color code indicated the identity of cells in E (E) DeltaF/F data for 10 example ROIs from D, and below lever position and reward times. (F) Average population activity of all SPNs aligned to cue presentation (light teal), rewarded lever press during cue blocks (dark teal), or rewarded lever presses during self-paced blocks (magenta). Shaded area is SEM. (G) Top: raster plots of grand average SPN responses (each row is a neuron), aligned to the cue presentation in cued blocks (left), the rewarded lever press in cued blocks (middle) and rewarded lever press in self-paced blocks (right). Cells are ordered by hierarchical clustering according to their average activity during the self-paced lever presses with cluster 1 at the top in blue, cluster 2 in the middle in red, and cluster 3 at the bottom in yellow. Bottom: Corresponding grand average z-scored deltaF/F response for each cluster, aligned to the cue or press, as in the rasters above. (H) Left: quantification of the peak deltaF/F z-scored response of cluster 1, aligned to the cue (light teal), the press in cued blocks (dark teal) and the press in self-evoked blocks (pink). The difference between these groups was not significant (One-way ANOVA p = 0.26 with Tukey-Kramer post-hoc test, ns p > 0.05). Middle: quantification of the peak deltaF/F z-scored response of cluster 2, aligned to the cue (light teal), press in cued blocks (dark teal) and press in self-evoked blocks (pink). This group showed the largest peak response to the cue in cued blocks, which was significantly larger than the peak response when aligned to the press in cued blocks or press in self-paced blocks (One-way ANOVA p < 1 x 10^-10^ with Tukey-Kramer post hoc test: ****p < .0001, ns p < 0.05). Right: quantification of the peak deltaF/F z-scored response of cluster 3, aligned to the cue (light teal), press in cued blocks (dark teal) and press in self-evoked blocks (pink). This group showed the largest peak response when aligned to the press in self-evoked blocks (One-way ANOVA p = 0.002 with Tukey-Kramer post hoc test: ****p < .0001, ns p > 0.05). Bar plots are mean ± sem.

Unsupervised clustering revealed that neurons (n = 1271) segregated into three functional ensembles, one of which—Cluster 1—was consistently active around lever pressing across both contexts. Notably, this shared movement-timed ensemble emerged spontaneously, underscoring that neurons encoding action are inherently shared across initiation modes. The activity of neurons belonging to each cluster can be visualized aligned either to the cue in cue-evoked blocks or to the press in both cue and self-paced blocks (Fig. 2G). Cluster 1 exhibited a peak response ∼200 ms after cue onset when aligned to it, which was also evident when aligned to the press (Fig. 2G, blue trace). There was no significant difference in peak amplitude of this cluster when aligned to the tone in cued blocks or to the press in either context (Fig. 2H, left; z-score responses: cued, tone-aligned = 0.49 ± .04; cued, press-aligned = 0.47 ± .04; self-paced, press-aligned = 0.56 ± .04; n = 118; one-way ANOVA, p = 0.27). This indicates that neurons in Cluster 1 are consistently active during lever pressing regardless of whether it is cue-evoked or self-initiated, revealing a shared neural representation of the action across behavioral contexts.

Neurons in Cluster 2 showed a grand average peak response well aligned with the cue, peaking ∼66 ms after tone onset. Their responses were significantly higher when aligned to the cue, indicating selective engagement by external sensory (Fig. 2H, middle, z-score responses: cued, tone aligned= 0.17 ± .004, cued, press aligned = 0.06 ± .008, self, press aligned = 0.06 ± .007, n = 868, One-way ANOVA p < 1 x 10^-10^). This was further confirmed by separating trials into bins based on reaction time (e.g., 0–333 ms or 334–666 ms; Fig. S2). In these trials, Cluster 2 neurons showed clear peaks at cue onset that preceded movement initiation (Fig. S2B, S2E).

Finally, Cluster 3 neurons peaked ∼1 s after cue onset, consistent with licking and/or reward consumption. They showed the strongest responses in self-paced trials when aligned to the press (Fig. 3H, right, z-score responses: cued, tone aligned= 0.34 ± .02, cued, press aligned = 0.39 ± .02, self, press aligned = 0.43 ± .02, n = 868, One-way ANOVA, p = 0.0015).

**Figure 3:**
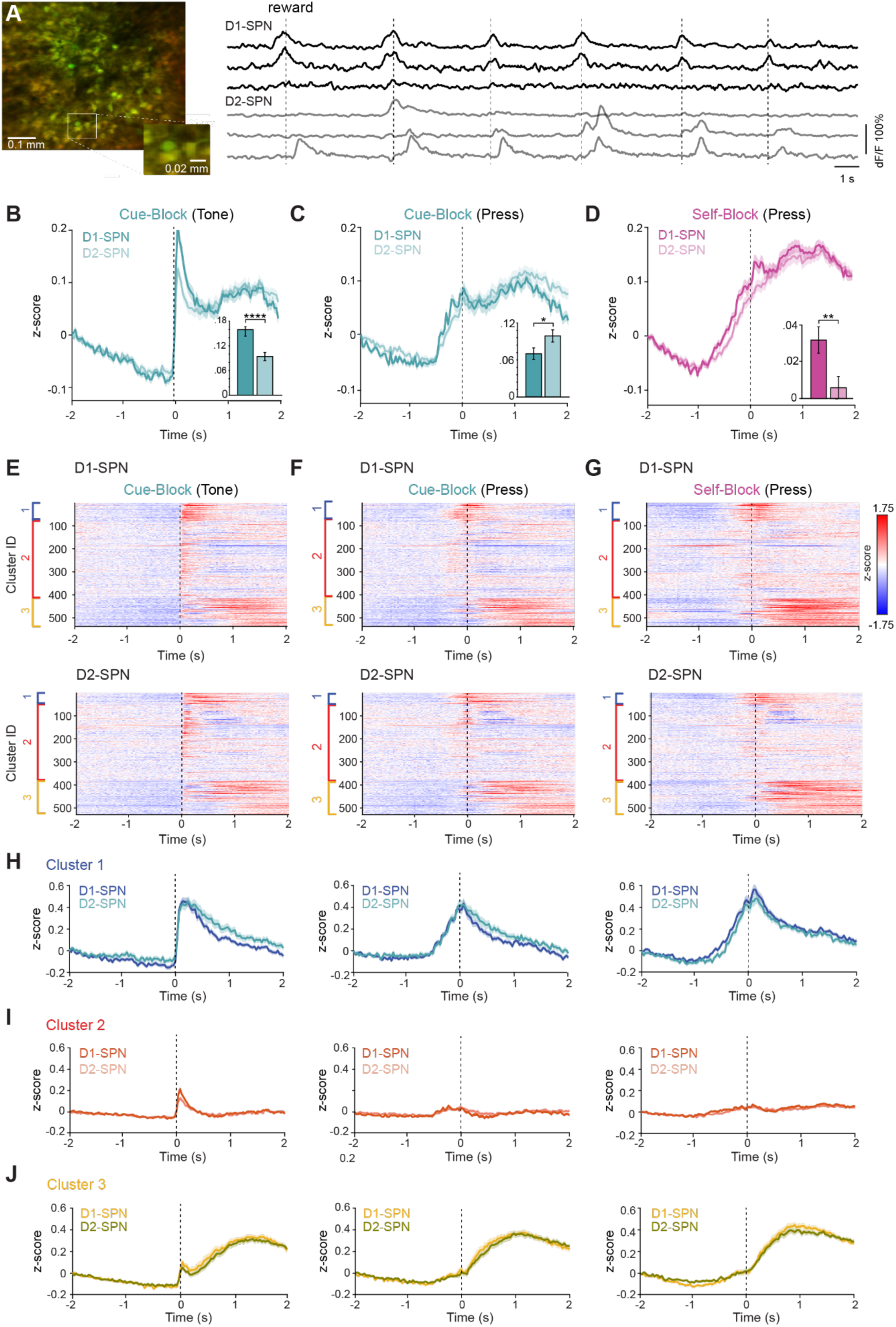
D1- and D2-SPN activity during the switching task. (A) Left: 2P image of FOV in an example animal showing expression of GCaMP8m (green) and TdTomato (red). Right: example deltaF/F from 3 putative D1-SPNs (black traces, top) and 3 D2-SPNs (grey traces, bottom). (B) Average population activity of D1-SPNs (dark teal) and D2-SPNs (light teal) aligned to cue presentation. Shaded area is SEM. Inset: Mean z-scored deltaF/F activity 0-200ms after the cue onset was significantly higher in D1-SPNs (Student’s t-test ****p < 0.0001). (C) Average population activity of D1-SPNs (dark teal) and D2-SPNs (light teal) aligned to lever press in cue blocks. Shaded area is SEM. Inset: Mean z-scored deltaF/F activity 1-2 s after the cue onset was significantly higher in D2-SPNs (Student’s t-test *p < 0.05). (D) Average population activity of D1-SPNs (dark magenta) and D2-SPNs (light magenta) aligned to reward lever press in self-evoked blocks. Shaded area is SEM. Inset: Mean z-scored deltaF/F activity -1s-0 s relative to cue onset was significantly higher in D1-SPNs (Student’s t-test **p < 0.01). (E) Raster plot of grand average D1-SPN responses (top) and D2-SPN responses (bottom), aligned to the cue presentation in cued blocks. Cells are ordered by cluster with cluster 1 at the top (blue), cluster 2 in the middle (red) and cluster 3 at the bottom (yellow). (F) As in (E), aligned to the rewarded lever press in cued blocks. (G) As in (E) and (F) aligned to the rewarded lever press in self-paced blocks. (H) Grand average z-scored deltaF/F traces for cluster 1, aligned to the cue in cue blocks (left), to the press in cue blocks (middle) or to the press in self-paced blocks (right). (I) As in H, for cluster 2. (J) As in H, for cluster 3.

Together, these results show that DLS neurons segregate into three functional ensembles—one encoding movement regardless of initiation mode, one selectively engaged by external cues, and one linked to post-action reward-related activity—highlighting distinct but complementary roles in processing motor and contextual information.

### D1- and D2-SPNs exhibit distinct temporal dynamics around action execution and post action periods

We used our ability to identify neurons as D1-SPNs or D2-SPNs by their expression of TdTomato and analyzed our data after separating ROIs into putative D1-SPNs or D2-SPNs (Fig. 3A). We then plotted the grand average z-scored fluorescence changes for all neurons imaged across the 6 mice (n = 537 D1-SPNs, n = 529 D2-SPNs). When aligned to the cue, D1-SPNs showed significantly higher responses in the first 200 ms after the cue (Fig. 3B; mean z-score 0-200 ms from cue onset: D1 = 0.16 ± 0.01, D2 = 0.09 ± 0.01, p = 2 x 10^-5^, Student’s t-test). D2-SPNs showed significantly higher average responses in the period from 1-2 s after the cue onset (Fig. 3C; mean z-score 1-2 s from cue onset: D1 = 0.07 ± 0.01, D2 = 0.10 ± 0.01, p = .03, Student’s t-test). D1-SPNs also showed significantly higher activity in the time period immediately preceding the press in both cue evoked and self-paced blocks (Fig. 3D; mean z-score -750-0 ms from cue onset aligned to press in self-paced block: D1 = 0.03 ± 0.007, D2 = 0.006 ± 0.006, p = 0.006, Student’s t-test).

Separating the previously clustered neurons into D1- and D2-SPNs revealed a similar proportion of each cluster across both cell types (Fig. S3). Cluster 1 neurons were active at cue onset and around the press in both trial types; within this cluster, D2-SPNs showed higher activation in the period 1-2 s after the cue, before the next intertrial interval (Fig. 3H, S3A–B).

Cluster 2 neurons were most strongly aligned to the cue, with D1-SPNs showing significantly higher responses in the 0–200 ms window after cue onset (Fig. 3I, left; Fig. S3D). Cluster 3 neurons peaked later, around reward and licking, and showed similar activation between D1-and D2-SPNs (Fig. 3J). These results reveal that differences between D1- and D2-SPNs are not homogeneous across the population, but emerge in specific functional clusters—D1-SPNs are more engaged during early cue processing and movement preparation in certain ensembles, whereas D2-SPNs show enhanced post-action activation in others—indicating that each pathway contributes to initiation and evaluation processes in a cluster- and context-dependent manner.

### Neural dynamics predict cue-evoked versus self-paced actions

The shared movement-timed neurons (Cluster 1) provided a foundation for testing whether population dynamics—rather than cell identity—differentiate internally and externally initiated actions. To do so, we examined features such as baseline activation levels, the temporal dynamics of activity buildup, and used a support vector machine (SVM) classifier to assess whether patterns of population activity could reliably predict trial type. These approaches allowed us to explore whether the mode of initiation is reflected not in the identity of responsive neurons, but in the structure or timing of their activity. Indeed, while many cells were responsive during both cued and self-paced lever presses, a substantial proportion exhibited differences in the amplitude of their responses across these trial types, suggesting that the strength of neural activation may encode information about the mode of action initiation (Fig. 4A). Principal component analysis of the population activity in an example mouse revealed that, although neural trajectories began in a similar state across conditions, they diverged during the preparatory period and then converged again at the time of reward delivery (Fig. 4B). This suggests that convergence at reward could reflect a shared outcome-related signal across initiation modes. Indeed, across all mice, the Euclidean distance between neural trajectories peaked several hundred milliseconds before the lever press, indicating a growing divergence in preparatory activity between cued and self-paced trials. This distance then dipped at the time of the press, reflecting transient convergence in neural dynamics associated with movement execution. Importantly, this was not a trivial consequence of differences in the overall averages, as subtraction of the grand average cue evoked and self-evoked traces did not show peaks either leading up to or following the press (Fig. 4C).

**Figure 4:**
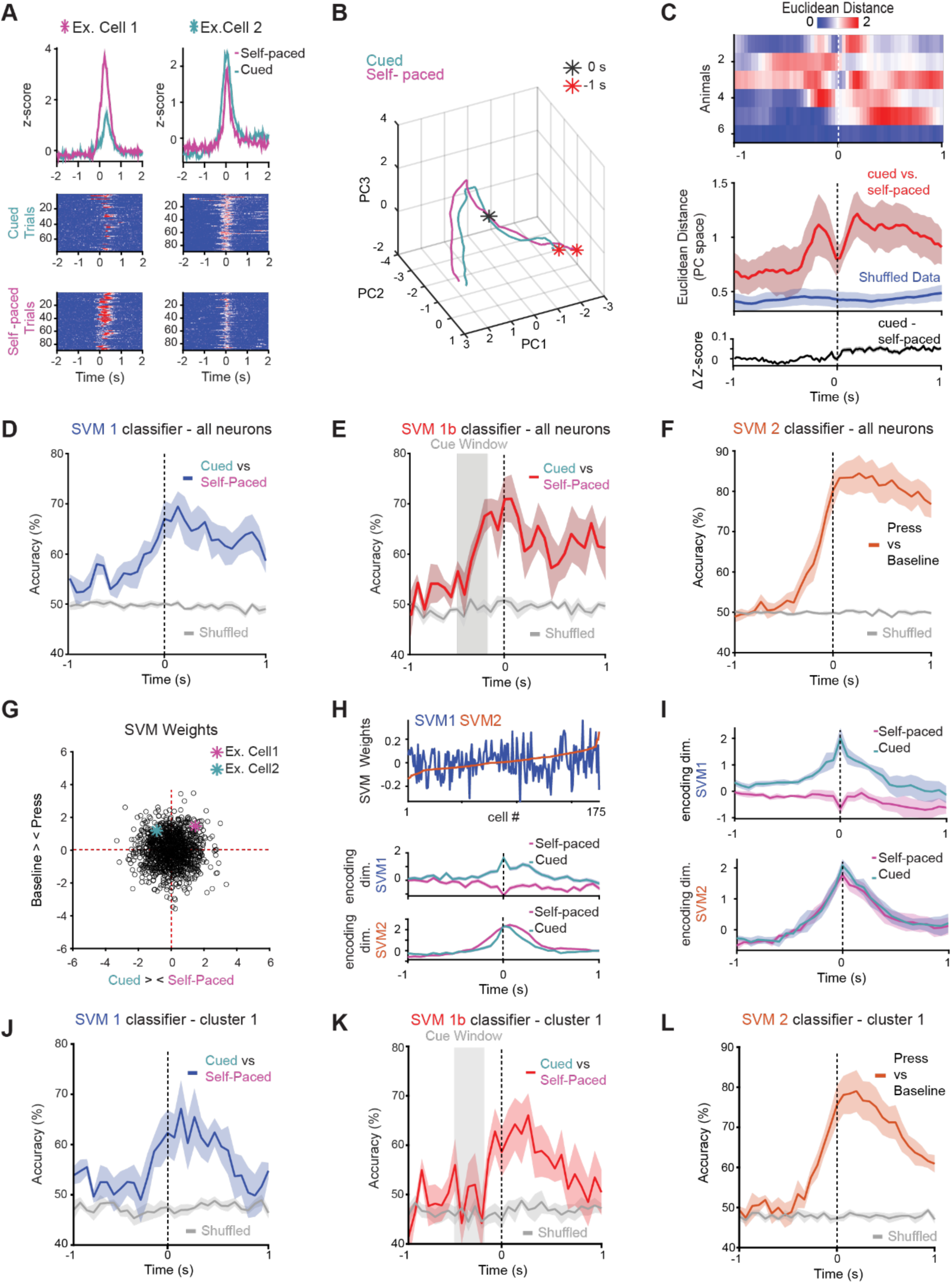
Distinct and Shared Neural Dimensions Underlying Self-Paced and Cue-Evoked Actions. (A) Example of two neurons that both show responses to cue evoked and self paced lever presses, albeit with different magnitudes. Grand average z-score responses plotted in top row, with raster plot heat maps of responses during cued and self-paced trials for each neuron underneath. (B) Principal component analysis (PCA) of population activity in an example mouse shows that neural trajectories for cue-evoked and self-paced trials. (C) Top: Euclidean distance between neural trajectories for cue-evoked and self-paced trials for individual mice. Middle: average Euclidean distance for all mice (red), compared to shuffled data (blue) showing that trajectories converge at the lever press and diverge during preparation and reward. The bottom panel shows the subtraction of grand-averaged ΔF/F lever-aligned traces, confirming that differences in trajectory are not explained by simple differences in mean activity between the two trial types. (D) Support vector machine classifier (SVM 1) trained on 90% of the data and tested on the remaining 10% successfully predicted whether lever presses were cue-evoked or self-initiated, indicating distinguishable population activity patterns across trial types. Gray plot is the prediction on shuffled data. (E) SVM classification (SVM 1b) accuracy rose above chance during the cue period (−495 to −165 ms before the press) for matched-latency trials. Gray plot is the prediction on shuffled data. (F) A classifier distinguishing lever presses from baseline (SVM2) performed above chance before movement onset, indicating that this ensemble contributes to a shared preparatory neural dimension across both action types. Gray plot is the prediction on shuffled data. (G) Plot of SVM weight distribution across all neurons, showing that it is broadly distributed. Neurons shown in panel A are highlighted. (H) Top: Neuron weights from SVM2 (press vs. baseline) plotted in order of magnitude (red), with overlaid weights from SVM1 (cue vs. self-paced) showing distinct, uncorrelated contributions to each classification axis. Population activity from an example mouse projected onto SVM-defined axes shows trial-type-specific separation peaking at movement onset (middle) and shared motor-related dynamics across conditions (bottom). (I) Same as H middle and bottom but for the entire population of neurons from all mice. (J) Same as D, for Cluster 1 neurons only. (K) Same as E, for Cluster 1 neurons only. (L) Same as F, for Cluster 1 neurons only. Shaded area in all plots is ± SEM

We next asked whether patterns of population activity contained information about the mode of initiation. To test this, we trained a support vector machine (SVM) classifier on 90% of the data and evaluated its performance on the remaining 10%, asking whether it could predict whether a given lever press was cue-evoked or self-initiated. The classifier performed above chance, with accuracy peaking at and just after movement onset (Fig. 4D), indicating that population dynamics around the time of execution contain features that distinguish initiation mode. When restricted to cued trials with similar reaction times, classification accuracy rose above chance even during the cue period, showing that predictive signals emerge before movement and likely reflect context-specific preparatory activity (Fig. 4E).

In parallel, we trained an SVM to discriminate lever presses—regardless of initiation context—from baseline. This classifier also performed above chance, beginning several hundred milliseconds before the press, revealing a shared, context-invariant preparatory dimension that generalized across action types (Fig. 4F). Thus, decoding analyses uncovered two complementary components of striatal activity: one encoding the context in which an action is generated, and another capturing a common motor execution signal that is conserved across contexts.

Examining classifier weights showed that discriminating information was distributed broadly across the population rather than concentrated in a small subset of cells (Fig. 4G). Moreover, neurons contributing strongly to the general “press vs. baseline” axis did not systematically contribute to the “self-paced vs. cue-evoked” axis, demonstrating that these context-invariant and context-specific signals occupy dissociable neural dimensions (Fig. 4H–I).

Analysis of SVM performance within functional clusters revealed that cluster 1 neurons—those with robust action-related responses shared across cue-evoked and self-paced trials—were surprisingly effective at distinguishing initiation mode (Fig. 4J). Despite showing highly similar press-aligned activity in both trial types, cluster 1 neurons supported above-chance classification of cue versus self-paced presses, performing as well as or better than cluster 2 (Fig. S4C-G), which contained cue-selective neurons. Notably, cluster 1 activity did not reliably encode trial type during the pre-press cue period (Fig. 4K), but it did predict lever pressing versus baseline, consistent with its role in shared motor preparation (Fig. 4L). This suggests that neurons that generalize across contexts at the single-cell level can nonetheless contribute to context-specific coding at the population level, highlighting how separable dimensions of activity within overlapping ensembles enable the striatum to flexibly represent both action execution and initiation mode.

Finally, when we performed PCA and trained SVM classifiers separately on D1- and D2-SPNs, both populations supported above-chance classification of trial type with similar accuracy. This indicates that context differentiation arises from population-level organization rather than cell-type segregation, despite distinct temporal profiles of D1- and D2-SPNs (Fig. S5).

## DISCUSSION

Our study reveals that the DLS encodes both externally triggered and internally generated actions through overlapping but distinguishable neural dynamics. Using a task that held motor output constant, we identified through unsupervised clustering a single population—Cluster 1—that consistently aligned with movement, regardless of initiation mode. We did not preselect these neurons; rather, they *emerged naturally* as the only action-timed ensemble, underscoring that neurons encoding action are inherently shared across contexts. Yet despite this overlap, the population dynamics within these shared neurons diverged prior to movement, reflecting distinct preparatory trajectories for cue-evoked versus self-initiated actions. This indicates that convergence between internal and external drivers does not occur only at the level of motor output (e.g., muscles or descending commands), but rather within **the striatal population space** itself.

The striatum thus forms a high-level “action space” where contextual information is integrated and transformed into a common execution signal—a context-generalizable code for action. In this view, flexibility in initiation arises not from switching between specialized circuits, but from dynamic reconfiguration within shared ensembles that can encode both context and action along orthogonal dimensions. This organization echoes emerging principles of population-level computation described in cortical circuits, where overlapping neurons support diverse functions through mixed selectivity and structured neural subspaces (*39–41*). In contrast to motor cortex, where preparatory activity converges across self-initiated and cue-driven movements (*6*), the striatum differentiates initiation context within shared ensembles, transforming convergent cortical plans into context-specific population dynamics.

Our ability to distinguish D1- and D2-SPNs further revealed distinct contributions of these two striatal subpopulations to context encoding. D1-SPNs were preferentially engaged during early cue processing and just prior to movement onset, consistent with a facilitative role during externally triggered actions. By contrast, D2-SPNs exhibited greater activity following lever presses, particularly in the post-cue and intertrial intervals, suggesting a role in outcome monitoring or behavioral suppression. These results refine classical models of direct and indirect pathways as opposing controllers of movement, showing instead that both contribute to shaping initiation and evaluation in a cluster- and context-dependent manner.

The simplicity of our behavioral paradigm is both a limitation and a strength. The one-dimensional lever press does not capture the richness of naturalistic motor sequences, but its stereotypy allows precise control over kinematics, isolating neural differences that reflect *initiation context* rather than movement variability. This design was essential for revealing that shared striatal populations flexibly encode both self-initiated and cue-triggered actions. Extending this framework to more complex, multidimensional behaviors will help determine whether the context- and action-related population subspaces identified here generalize across the broader motor repertoire.

## Materials and methods

### Animals

All experiments were carried out with 8-16-week-old male and female mice > 15 grams in accordance with National Institutes of Health (NIH) guidelines and protocols approved by the NYU Langone Health (NYULH) Institutional Animal Care and Use Committee (protocol# PROTO201900059). No effects of sex are reported. Mice were bred in-house on a C57Bl/6J background and housed in a reverse light/dark cycle (light 11 pm to 11 am), at a temperature of 22 ± 2°C with food available ad libitum. Mice were group housed with littermates of the same sex during experimental procedures. The following strains used in this proposal were sourced from Jackson Laboratories: Ai14: B6.Cg-*Gt(ROSA)26Sor^tm14(CAG-tdTomato)Hze^*/J (JAX:007914). The following bacterial artificial chromosome (BAC) transgenic mouse strain used in this paper were sourced from the Mutant Mouse Regional Resource Centers (MMRRC): D2-Cre: B6.FVB(Cg)-Tg(Drd2-cre)ER44Gsat/Mmucd (MMRRC_032108-UCD); D1-Cre: Tg(Drd1-cre)EY217Gsat/Mmucd (MMRRC_030778-UCD).

All experiments were carried out with 8-16-week-old male and female mice in accordance with protocols approved by the NYU Langone Health (NYULH) Institutional Animal Care and Use Committee (protocol #PROTO201900059). D2-Cre bacterial artificial chromosome (BAC) transgenic mice were obtained from Gene Expression Nervous System Atlas (GENSAT; founder line EY217 for D1-Cre, ER44 for D2-Cre), and purchased through the Mutant Mouse Regional Resource Centers (MMRRC). These mice were crossed with Lox-Stop-Lox-tdTomato mice (Ai14, JAX:007914).

The mice were housed in a reverse light/dark cycle (light 11 pm to 11 am), at a temperature of 22 ± 2°C with food restriction beginning 24-48 hours before the commencement of behavioral training.

### General surgical procedures

Mice were anesthetized with isoflurane (2.5%–3%, plus oxygen at 1–1.5 l/min) and then placed in a stereotaxic holder (Kopf Instruments). Meloxicam (5 mg/ml) was administered subcutaneously for analgesia prior to the onset of surgery. The scalp was shaved and disinfected using 70% alcohol followed by iodine, and intradermal Bupivacaine (2 mg/kg) was also provided for local anesthesia. The mouse’s body temperature was maintained throughout surgery at 37 °C using an animal temperature controller (FHC DC Temperature Controller). After surgery mice were allowed to recover until ambulatory in the home cage on a heating pad.

Additional analgesia was administered with ibuprofen (100 mg/ml) in the drinking water for 3 days post-surgery.

### Chronic GRIN lens implantation

Animals were unilaterally injected with 500 nl of AAV.CamKIIa.jGCaMP8m.WPRE (Addgene ref# 176751-AAV9, lot # v159649, 2.7E13 vg/ml), or AAV.syn.jGCaMP8f.WPRE (Addgene ref# 162376-AAV9, lot # v136270 ,2.3E13GC/ml). Injections were performed in the DLS (coordinates: AP +0.5 mm ML -2.2 mm and DV -2.6 mm from bregma). Tissue was then aspirated at a circumference of ∼ 1mm to a depth ∼200 µm above the center of the injection site. A 1.0 mm gradient index (GRIN) lens (length: ∼4.38 mm, working distance: image side: 0.1 mm in air, object side: 0.25 mm in water, design wavelength: 860 nm, NA 0.5, non-coated, catalog #1050-006242, Inscopix Inc) was then lowered ∼2mm in depth, stopping ∼200-400 μm above the center of the injection site. The lens was secured in place with glue and a headpost was implanted with C&B Metabond dental cement (Parkell) around the lens.

### Behavioral training and monitoring

Mice were subjected to a food restriction schedule, and were maintained at 85% of their starting weight. We implemented a custom head restrained lever-based training paradigm where mice push the lever spontaneously (“self-initiated” behavior) or push the lever in response to an auditory cue (“cued” behavior) to collect a to collect 5 µl of a 10% sucrose solution reward. We used an adapted custom designed lever(*42*), that was mounted onto a rotary encoder (US Digital) X cm from the lever handle. A magnet (CMS magnets) was mounted to the bottom of the lever and positioned 1.5 cm above a static magnet that established the resting position of the lever and provided a way to alter the movement resistance. The lever handle was positioned adjacent to a cup (copyright IR CU21353, 3D printed using a Makerbot Replicator+ 3D Printer) to hold mice below two plate clamps for head fixation.

The task required the mice to push the lever across a threshold of 5 mm of displacement to receive a reward. In self-initiated blocks, mice could get a reward when they spontaneously pushed the lever past the reward threshold. However, an “intertrial interval” was superimposed such that after a reward was given, 2-3 s needed to pass before another reward could be delivered. In addition, to discourage the mice from continuously pushing the lever, mice had to hold the lever at the home position (“quiet period”) for at least 1-2 s in before a rewarded lever press. In cued blocks, the intertrial interval was 4-8 s, after which, if they mice held the lever in the home position for a period of 1.5-2.5 s a ‘go’ cue (5 kHz pure tone) would be delivered. If mice pushed the lever past the reward threshold within 2 s after this cue, they would receive the reward. Training consisted in 3 stages. First, mice were rewarded for spontaneously lever pushes for 3 days of training. During this phase the ITI was held at 2-3s and the mice had to hold the lever still for at least 1-2 seconds between trials. Next, mice were trained to push the lever in response to the auditory cue, achieving a > 80% “hit” rate over a period of 5-7 training days. Finally, on the day of the recordings, mice alternated between cued and self-paced blocks, with each block lasting for a total of ∼30 rewards.

### Two-photon calcium imaging

All imaging experiments were conducted on a Ultima 2Pplus two-photon laser-scanning microscope (Bruker, Inc.). The system was configured with 8 kHz resonant-galvo-galvo laser scanning mirrors and imaging frames of 512 × 512 pixels (corresponding to an area of 1000 µm x 1000 µm) were acquired at 30 fps. The system was equipped with two-channel fluorescence detection with amplified non-cooled GAsP photomultiplier tubes (PMTs). Emitted fluorescence was first directed to the PMTs, then split into “green” and “red” channels by a 565 nm sharp edge long-pass dichroic mirror. The green channel and red channel were subsequently filtered by a 525 nm/39 nm bandpass filter, and 593 nm /40 nm bandpass filter, respectively, before detection in the PMTs. The microscope was controlled via Prairie View version 5.6.

Imaging was performed via a Nikon 16x, 0.8 NA water immersion objective placed over the implanted GRIN lens. A Coherent Chameleon Vision II tunable titanium-sapphire laser tuned to 920 nm with 75 fs pulses at 80 MHz repetition rate was used. Dispersion correction was adjusted to maximize fluorescent brightness as recorded under the objective. The imaging power was modulated through a Conoptics 350-105 Pockels Cell driven by a Conoptics 302 RM Amplifier.

Behavioral data including lever presses and video data was recorded using HARP Behavior board (https://open-ephys.org/harp/oeps-1216) and custom workflows using the BONSAI (https://bonsai-rx.org/) software package. Behavioral and imaging data was synchronized by sending TTL pulse from the 2P-microscope to the HARP behavior board every time a new 2P frame was recorded. These timestamps could then be used to align and down sample behavioral data to match imaging data using custom MATLAB scripts.

### Muscimol infusions

Adult mice (C57BL/6J, male or female, aged 8–12 weeks) were implanted with unilateral guide cannulae targeting the dorsal striatum. Mice were anesthetized and prepared for surgery as described above. A small craniotomy was made at the target coordinate for the dorsolateral striatum (anteroposterior: +0.5 mm, mediolateral: ±2.2 mm from bregma), or the posterior “tail” striatum (anteroposterior: -1.6 mm, mediolateral: ±3.1 mm from bregma) and a 26-gauge stainless steel guide cannula (Plastics One) was lowered to 0.5 mm above the final infusion site (dorsoventral: −2.15 mm for DLS, -2.75 for posterior striatum from the skull surface). The cannula was secured to the skull using dental acrylic (C&B Metabond). A dummy cannula was inserted to prevent occlusion. Mice were allowed to recover for at least 7 days before beginning behavioral or infusion experiments. Post-operative analgesia (meloxicam, 5 mg/kg, s.c.) was administered once daily for 3 days.

For acute inactivation of striatal activity, fluorescent muscimol (Muscimol, BODIPY™ TMR-X Conjugate; 0.5mg , ml^-1^ in sterile phosphate-buffered saline [PBS]) was infused unilaterally through an internal injector extending 0.5 mm beyond the guide cannula tip. Mice were gently head-fixed for infusion procedures.

A total volume of 350 nl fluorescent muscimol was delivered at a rate of 0.1 μL/min using a microsyringe pump (Hamilton). After infusion, the injector was left in place for an additional 1 minutes to allow diffusion and minimize backflow. The injector was then removed and replaced with the dummy cannula. Control animals received equal-volume infusions of sterile PBS.

Behavioral testing commenced 30 minutes after infusion to ensure effective onset of muscimol action. The location of the infusion site was later verified histologically.

### Immunohistochemistry

Mice were deeply anesthetized with isoflurane (5 % for 5-8 min, inhaled) and perfused transcardially with 4 % paraformaldehyde (PFA) in 0.1 M phosphate buffered saline (PBS). Brains were removed and fixed in 4 % PFA for a maximum of 24 hours in the same solution, which was then replaced by a 0.1 M PBS solution 100 µm coronal slices were cut (vibratome, Leica VT1000S). The slices were subsequently mounted with DAPI mounting media (Southern Biotech). Images were obtained with an Olympus VS2000) slide scanner.

## Quantification and statistical analysis

### Calcium imaging data preprocessing

Calcium imaging data was motion corrected using non-rigid motion correction, cells were segmented and raw fluorescence traces extracted using Suite2p (0.10.0)^5,6^. Automatically classified cells were further manually curated using the Suite2p GUI and occasional ROIs with clearly non-physiological morphological features (order of magnitude to large/small), no transients or clearly non-physiological activity patterns (highly regular or square shaped fluctuations), were excluded.

Using custom MATLAB code, raw fluorescence traces were neuropil-corrected by subtracting 0.7 times the neuropil signal from the raw cellular fluorescence. For each neuron, the fluorescence signal was then detrended to compute ΔF/F. This was done by estimating the baseline activity for each cell by applying gaussian smoothing followed by a 50th percentile filter within non-overlapping 60-second windows across the full recording. The resulting baseline trace was subtracted from the raw signal and divided by the same baseline to yield a ΔF/F signal, expressed as a percentage. Any traces with negative baseline values were excluded.

After baseline correction, each ΔF/F trace was z-scored across the entire imaging session. These z-scored traces were then used for all subsequent analyses, including trial alignment, averaging, and dimensionality reduction.

### Identification of D1- and D2-SPNs

To classify individual neurons as either direct pathway spiny projection neurons (D1-SPNs) or indirect pathway spiny projection neurons (D2-SPNs), we used a custom MATLAB graphical user interface (GUI) to manually inspect each region of interest (ROI). Classification was based on the spatial correspondence between GCaMP-expressing cell bodies and tdTomato fluorescence. The experimenter was blinded to the experimental condition during this process. In EY217xAi14 mice, ROIs were classified as putative D1-SPNs if a tdTomato-positive signal spatially overlapped with the GCaMP signal and the shape of the labeled soma was clearly distinguishable from the background. In contrast, ROIs lacking a tdTomato signal, or showing a non-overlapping tdTomato shape, were classified as putative D2-SPNs. For D2-Cre x Ai14 mice, the criteria were reversed: overlapping GCaMP and tdTomato signals indicated putative D2-SPNs, while tdTomato-negative or mismatched cells were categorized as putative D1-SPNs. ROIs that could not be reliably classified based on these criteria were excluded from further D1-/D2-SPN-specific analyses.

### Population activity analyses

To compute peri-event trial-averaged activity, the neural activity of individual neurons was aligned to task-relevant events (i.e., cue onset or lever threshold crossing) and averaged across trials. For each neuron, calcium traces were first z-scored, then segmented into peri-event windows (−5 to +5 seconds, sampled at 30 Hz) around each event. Trials contaminated by non-physiologically large artifacts were excluded using amplitude thresholds. For each event type (cued, uncued, or cue-only), the trial-averaged activity of each neuron was computed by taking the mean trace across all valid trials. To compare average population activity across task blocks, the trial-averaged activity of all cells from all animals was pooled and averaged.

### Hierarchical clustering

To identify functional subpopulations of neurons, we performed hierarchical clustering on the trial-averaged activity of all recorded neurons (n = 1271) aligned to the cue in cue-evoked trials. Each neuron’s response was defined as its z-scored calcium trace in a time window spanning 4 seconds around the cue (−2 to +2 s). We computed the pairwise Euclidean distance between these activity traces and applied Ward’s method for agglomerative hierarchical clustering.

Neurons were grouped into three clusters by cutting the resulting dendrogram at a fixed number of clusters (k = 3). The resulting cluster assignments were used to sort neurons for subsequent visualization across task conditions.

### Principal component and Population Rate Vector Analysis

For population trajectory analysis, we categorized trials into self-paced (uncued) and externally triggered (cued) actions and computed the trial-averaged trajectory in the space of the first three principal components (PCs) for each condition. Euclidean distance between the trajectories was calculated at each time point to quantify their divergence over time. As a control, we repeated the analysis 100 times with randomly shuffled trial labels to generate a null distribution of distances and compute the mean and standard error of the shuffled results.

### Support Vector Machine

To classify cue-based versus self-initiated lever presses using neuronal activity, we trained a linear Support Vector Machine (SVM) classifier on calcium imaging data from individual animal sessions. Equal numbers of trials from each condition were randomly sub-sampled, and for each trial, fluorescence signals were aligned to behavioral events and segmented into 66 ms bins (2 imaging frames per bin) across a 4-second peri-event window. At each time bin, the SVM was trained on the average binned activity across neurons to distinguish between trial types, allowing us to track decoding accuracy over time. Classification was performed using 10-fold cross-validation. This procedure was repeated across all time bins to obtain a time-resolved decoding performance curve, with the resulting accuracies averaged across folds and sessions for further analysis. To estimate chance-level performance, we randomly shuffled the condition labels (cue-evoked vs. self-initiated) across trials while keeping the underlying calcium activity unchanged. We then retrained the SVM using the same cross-validation procedure, repeating this process multiple times to generate a null distribution of decoding accuracies for comparison with the real data.

### Statistical analysis

Data are presented as mean ± SEM. Statistical analyses were performed using MATLAB custom code. For unpaired datasets, two-tailed Student’s *T*-tests (for normally distributed datasets) or Mann–Whitney tests (for nonnormally distributed datasets) were employed. For paired datasets, two-tailed Student’s Paired *T*-test or Wilcoxon Signed-Rank test (for nonnormally distributed datasets) were employed. For multiple comparisons, we used Two-Way ANOVA followed by Sidak’s test. Values of *P* < 0.05 were considered statistically significant. *P* values are reported as follows: \**P* < 0.05; \*\**P* < 0.01; \*\*\**P* < 0.001; \*\*\*\**P* < 0.0001. In all plots, unless otherwise noted, errors are plotted as ±SEM.

## Acknowledgements

We thank Karin Morandell and David Schneider for help with implementation of the lever, Hélio Rodrigues for the design and build of the behavioral boxes we used for mouse training, and Corryn Chaimowitz for technical assistance.

## Author contributions

T.S., R.C. and V.R.A. contributed to the study design and plan for data analysis. S.F.P, J.K, S.S and I.R.V performed all experiments. J.K and S.F.P created a database for all in vivo data and wrote the custom code to analyze the data and make the figures. T.S., R.M.C and V.R.A. wrote the paper.

