## Supplementary Figures S1-S5 for "Shared striatal neurons exhibit context-specific dynamics for internally and externally driven actions"

### Supplementary Figures and Legends

#### Figure S1

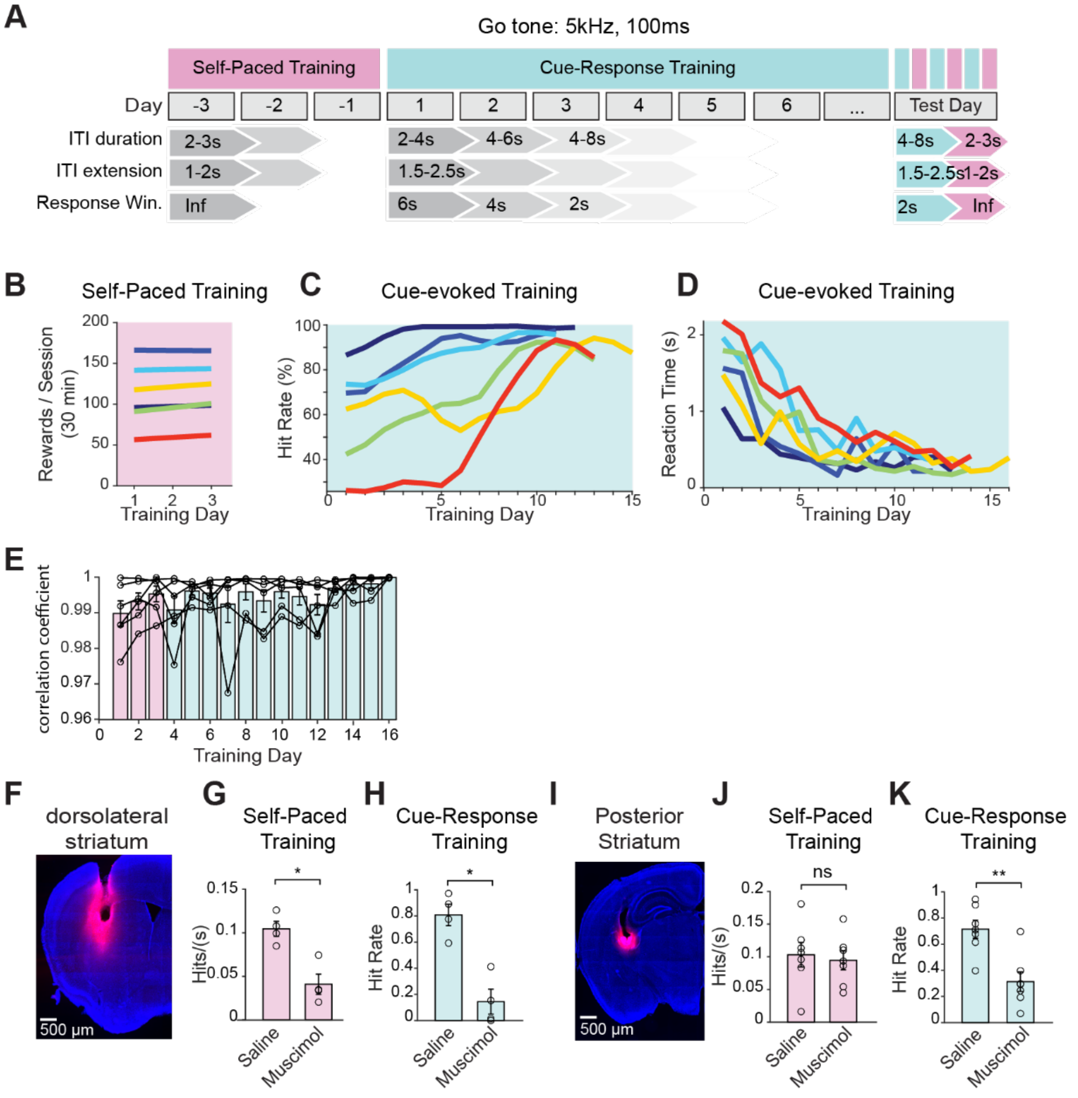


**Figure S1**, related to Figure 1: Learning of the cue-evoked/self-paced lever switching task

(A) Schematic of the training paradigm and timeline for acquisition of the switching task.
(B) The number of rewards per session during self-paced training, plotted for all 6 mice.
(C) The performance (plotted as hit rate) on each day cue-evoked training of all 6 mice.
(D) The reaction time on each day of cue-evoked training of all 6 mice.
(E) Average lever press trajectory correlation coefficient across all training sessions for each individual animal, showing that lever presses become more correlated (stereotypical) over time. Bars are mean ± sem.
(F) Histological image showing muscimol injection in the dorsolateral striatum.
(G) Hit rate for self-paced training was significantly reduced by muscimol injection in the DLS (saline: 0.10 ± 0.009 hits/s; muscimol 0.04 ± 0.01 hits/s; n = 4, Mann-Whitney U test: $*p < 0.05$). Bar plots are mean ± sem.

(H) Hit rate during cue-evoked training was significantly affected by muscimol injection in the "anterior" striatum, or DLS (saline: 0.81 ± 0.08; muscimol 0.19 ± 0.10 hits/s; n = 4, Mann-Whitney U test: *$p < 0.05$). Bar plots are mean ± sem.

(I) Histological image showing muscimol injection in the posterior striatum.

(J) Hit rate for self-paced training was not affected by muscimol injection in the posterior, or "tail" of the striatum (saline: 0.10 ± 0.02 hits/s; muscimol 0.09 ± 0.01 hits/s, n = 7, Mann-Whitney U test: ns $p > 0.05$). Bar plots are mean ± sem.

(K) Hit rate during cue-evoked training was significantly affected by muscimol injection in the posterior striatum (saline: 0.71 ± 0.07; muscimol 0.31 ± 0.07 hits/s; n = 7, Mann-Whitney U test: **$p < 0.01$).

Bar plots are mean ± sem.

Klee, Fernando-Peiris et al.

#### Figure S2

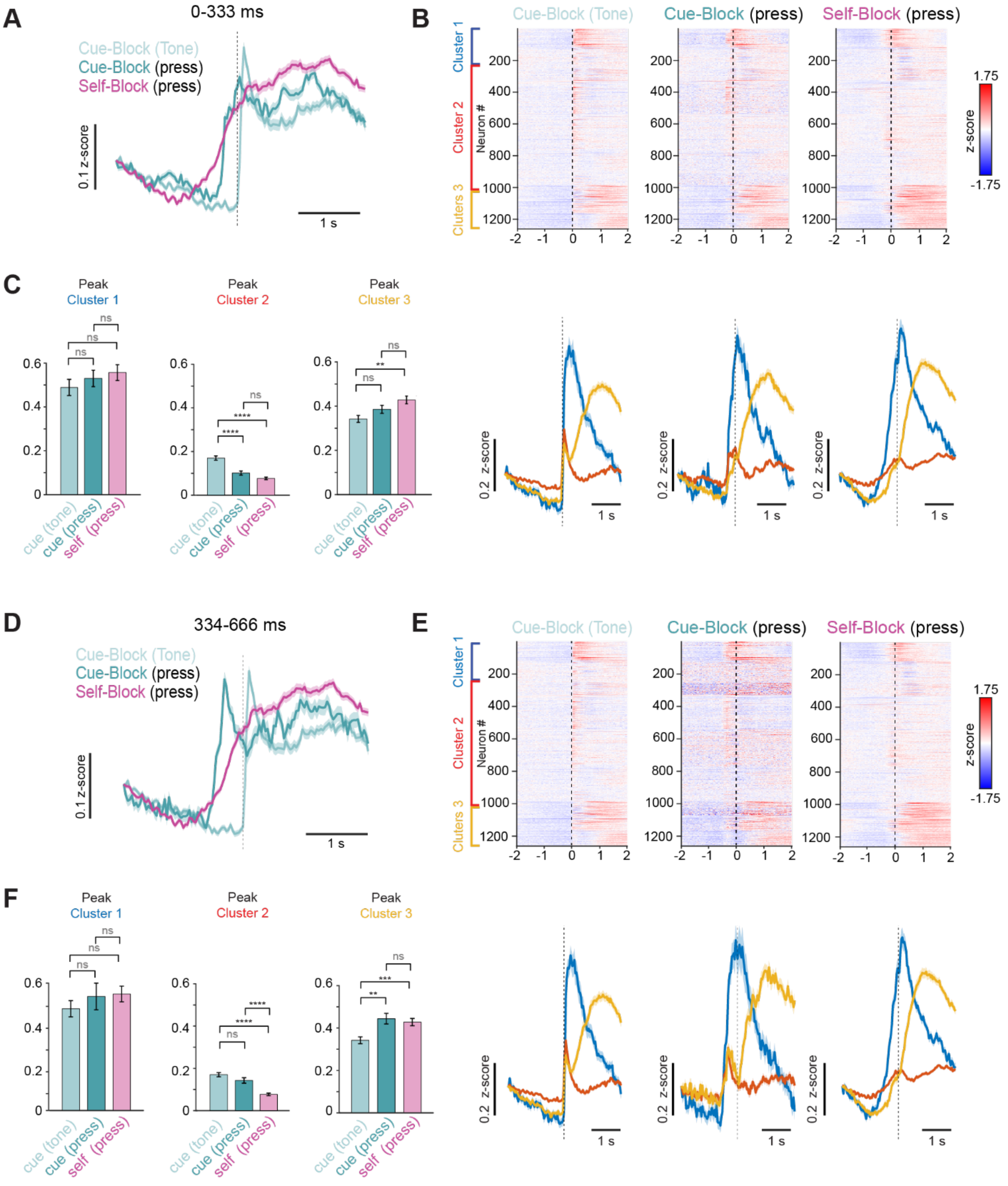


**Figure S2**, related to Figure 2: Separation of the trials by response time reveals a distinct cue response in cluster 2 neurons

(A) Grand average z-scored delta F/F responses of all SPNs for trials with reaction times between 0–333 ms) for cued blocks aligned to the tone (light teal), cued blocks aligned to the press (dark teal) and self-paced blocks (magenta).
(B) Top: Raster plots of z-scored delta F/F responses of all neurons for trials with 0–333 ms reaction times, organized by cluster with cluster 1 (blue) on top, cluster 2 (red) in the middle and cluster 3 (yellow) at the bottom. Left is aligned to cue onset in cued blocks, middle is aligned to lever press in cued blocks and right is aligned to lever press in self-paced blocks. Bottom: Corresponding grand average z-scored deltaF/F response for each cluster, aligned to the cue or press, as in the rasters above.
(C) Left: quantification of the peak deltaF/F z-scored response of cluster 1, aligned to the cue (light teal), the press in cued blocks (dark teal) and the press in self-evoked blocks (pink). The difference between these groups was not significant (z-scored peak responses: cue aligned = 0.49 ± 0.04, cued, press aligned = 0.53 ± 0.04, self, press aligned = 0.56 ± 0.04, n = 1271, One-way ANOVA $p = 0.46$ with Tukey-Kramer post-hoc test, ns $p > 0.05$). Middle: quantification of the peak deltaF/F z-scored response of cluster 2, aligned to the cue (light teal), press in cued blocks (dark teal) and press in self-evoked blocks (pink). This group showed the largest peak response to the cue and press in cued trials, which were both significantly larger than the peak response in the self-paced trials (z-scored peak responses: cue aligned = 0.17 ± 0.01, cued, press aligned = 0.10 ± 0.01, self, press aligned = 0.08 ± 0.01, n = 1271, One-way ANOVA $p < 1 \times 10^{-10}$ with Tukey-Kramer post hoc test: ****$p < .0001$, ns $p > 0.05$). Right: quantification of the peak deltaF/F z-scored response of cluster 3, aligned to the cue (light teal), press in cued blocks (dark teal) and press in self-evoked blocks (pink). This group showed the the largest peak response when aligned to the press in both cued and self-paced blocks (z-scored peak responses: cue aligned = 0.34 ± 0.02, cued, press aligned = 0.39 ± 0.02, self, press aligned = 0.43 ± 0.02, n = 1271, One-way ANOVA $p = 0.002$ with Tukey-Kramer post hoc test: **$p < 0.01$, ns $p > 0.05$). Bar plots are mean ± sem.
(D) As in (A) but for trials in which reaction times were between 334 – 666 ms.
(E) As in B, but for trials in which reaction times were between 334 – 666 ms.
(F) Left: quantification of the peak deltaF/F z-scored response of cluster 1, aligned to the cue (light teal), press in cued blocks (dark teal) and press in self-evoked blocks (pink). The difference between these groups was not significant (z-scored peak responses: cue aligned = 0.49 ± 0.04, cued, press aligned = 0.55 ± 0.06, self, press aligned = 0.56 ± 0.04, n = 1271, One-way ANOVA $p = 0.53$ with Tukey-Kramer post-hoc test, ns $p > 0.05$). Middle: quantification of the peak deltaF/F z-scored response of cluster 2, aligned to the cue (light teal), press in cued blocks (dark teal) and press in self-evoked blocks (pink). This group again showed the largest peak response to the cue and press in cued trials, which were significantly larger than the peak response in the self-paced trials (z-scored peak responses: cue aligned = 0.17 ± 0.01, cued, press aligned = 0.14 ± 0.01, self, press aligned = 0.08 ± 0.01, n = 1271, One-way ANOVA $p = 1.3 \times 10\text{-}^{10}$ with Tukey-Kramer post hoc test: ****$p < .0001$, ns $p > 0.05$). Right: Middle: quantification of the peak deltaF/F z-scored response of cluster 3, aligned to the cue (light teal), press in cued blocks (dark teal) and press in self-evoked blocks (pink). This group showed the largest peak response to the press in cued and self-paced trials, which were significantly larger than the peak response in the cue aligned trials (z-scored peak responses: cue aligned = 0.34 ± 0.02, cued, press aligned = 0.44 ± 0.02, self, press aligned = 0.43 ±

0.02, n = 1271, One-way ANOVA p = 0.005 with Tukey-Kramer post hoc test: **p < 0.01, ***p < 0.001, ns p > 0.05).

Bar plots are mean ± sem.

**Figure S3**

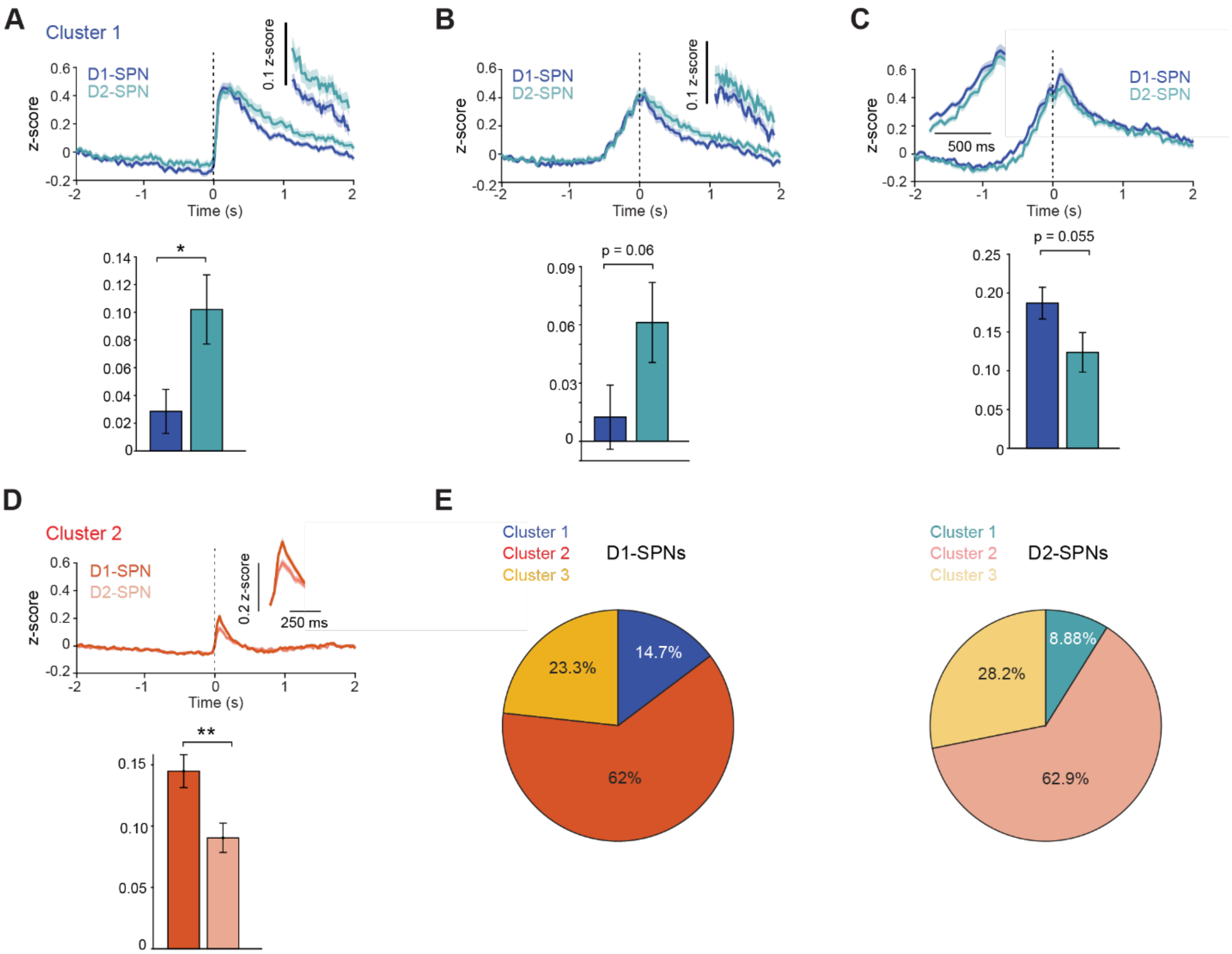


**Figure S3**, related to Figure 3: Quantification of average D1 and D2 responses by cluster reveals larger cue-evoked activation in cluster 2 D1-SPNs

(A) Responses of cluster 1 D1-SPNs (dark blue) and D2-SPNs (teal) aligned to the tone in the cued blocks with inset showing area quantified in bar graph below (1-2 s post press). D2-SPNs in this cluster showed a significantly larger response than D1-SPNs in this time window (mean z-score 1 -2 s from cue onset aligned to press in self-paced block: D1 = 0.03 ± 0.016, D2 = 0.10 ± 0.02, Student's t-test, *p < 0.05). Bar plots are data ± SEM.

(B) Responses of cluster 1 D1-SPNs (dark blue) and D2-SPNs (teal) aligned to the press in the cued blocks with inset showing area quantified in bar graph below, between 1 and 2 s post press. D2-SPNs in this cluster showed a higher response than D1-SPNs in this time window, although this difference was not significant (mean z-score 1 -2 s from cue onset aligned to press in self paced block: D1 = 0.012 ± 0.017, D2 = 0.06 ± 0.02, Student's t-test, $p = 0.06$). Bar plots are data ± SEM.

(C) Responses of cluster 1 D1-SPNs (dark blue) and D2-SPNs (teal) aligned to the press in the cued blocks with inset showing area quantified in bar graph below, between -750 – 0 ms before the press. D1-SPNs in this cluster showed a higher response than D2-SPNs in this time window, although this difference was not significant (mean z-score -750 – 0 ms from cue onset aligned to press in self paced block: D1 = 0.19 ± 0.02, D2 = 0.12 ± 0.03, $p = 0.05$, Student's t-test). Bar plots are data ± SEM.

(D) Responses of cluster 1 D1-SPNs (dark red) and D2-SPNs (light red) aligned to the tone in the cued blocks with inset showing area quantified in bar graph below, between 0 – 200 ms after the tone onset. D1-SPNs in this cluster showed a significantly higher response than D2-SPNs in this time window (mean z-score 0 – 200 ms from cue onset aligned to press in self paced block: D1 = 0.14 ± 0.01, D2 = 0.09 ± 0.01, Student's t-test, $^{**}p < 0.01$). Bar plots are data ± SEM.

(E) Pie charts showing proportion of cluster 1, 2 and 3 neurons in D1-SPN (left, darker colors) and D2-SPN (right, lighter colors) subpopulations.

Shaded area in all line plots is ± SEM

#### Figure S4

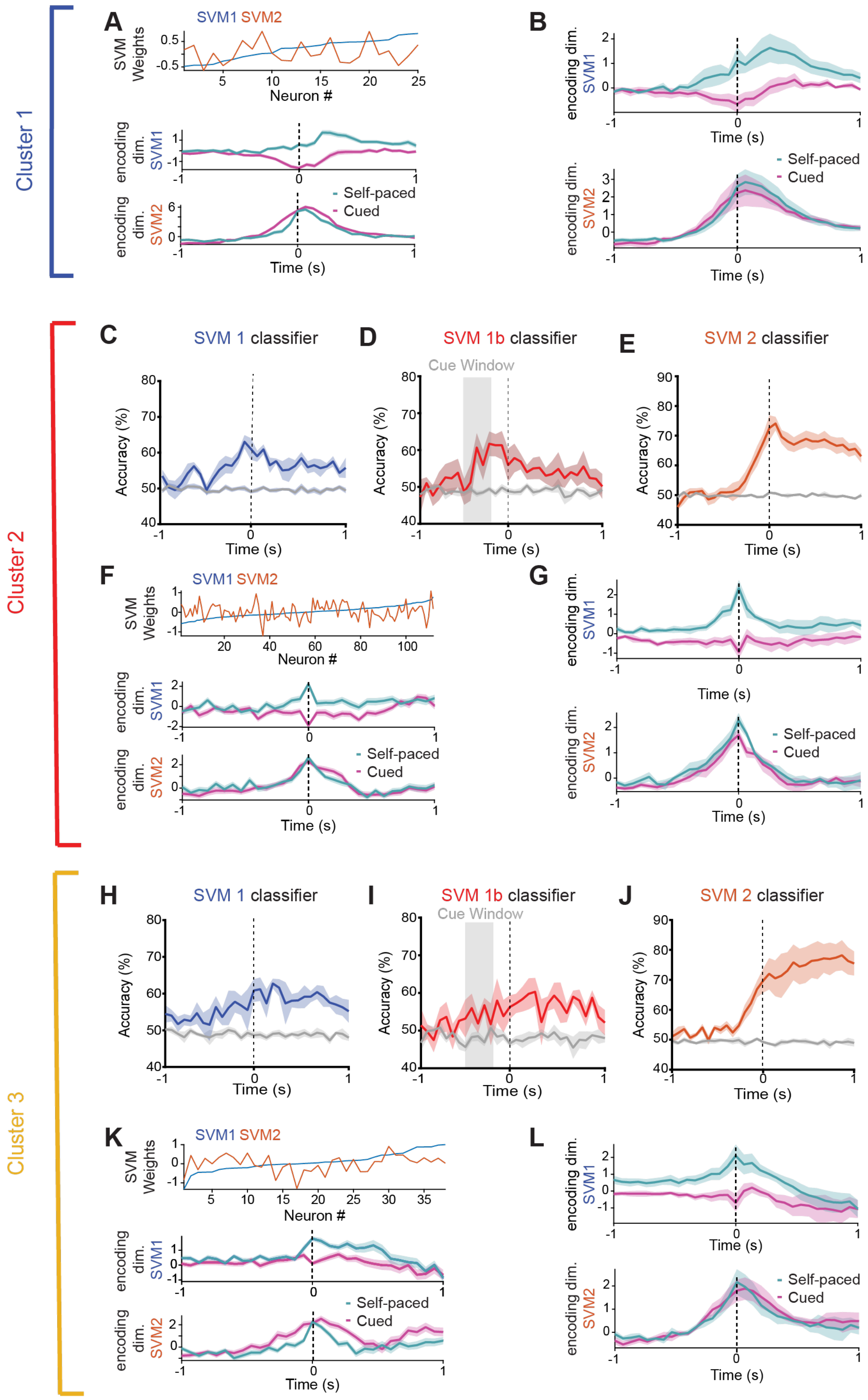
Cluster 1
Cluster 2
Cluster 3
A
B
C
D
E
F
G
H
I
J
K
L
SVM1 SVM2
SVM Weights
Neuron #
encoding dim. SVM1
encoding dim. SVM2
Self-paced
Cued
Time (s)
SVM 1 classifier
SVM 1b classifier
SVM 2 classifier
Cue Window
Accuracy (%)

**Figure S4**, related to figure 4, SVM of neuronal data by cluster reveals shared action-related cells in cluster 1 action specific neurons.

(A) For Cluster 1 neurons, Top: Neuron weights from SVM2 (press vs. baseline) plotted in order of magnitude (red), with overlaid weights from SVM1 (cue vs. self-paced) showing distinct, uncorrelated contributions to each classification axis. Population activity from an example mouse projected onto SVM-defined axes shows trial-type-specific separation peaking at movement onset (middle) and shared motor-related dynamics across conditions (bottom).
(B) Same as A middle and bottom, for all Cluster 1 neurons
(C) For cluster 2 neurons a support vector machine (SVM 1) classifier trained on 90% of the data and tested on the remaining 10% successfully predicted whether lever presses were cue-evoked or self-initiated based on activity, indicating that this subpopulation carries distinguishable population activity patterns across trial types. Gray plot is the prediction on shuffled data.
(D) For cluster 2 neurons, SVM classification accuracy rose above chance during the cue period (−495 to −165 ms before the press, SVM 1b) in matched-latency trials, indicating that this subpopulation reliably encoded movement type prior to lever press onset. Gray plot is the prediction on shuffled data.
(E) For cluster 2 neurons, the classifier distinguishing lever presses from baseline (SVM2) performed above chance before movement onset, indicating that this ensemble contributes to a shared preparatory neural dimension across both action types. Gray plot is the prediction on shuffled data.
(F) Same as (A), for cluster 2 neurons
(G) Same as (B), for cluster 2 neurons
(H) Same as (C), for cluster 3 neurons.
(I) Same as (D), for cluster 3 neurons, For cluster 3 neurons, SVM classification accuracy did not rise above chance during the cue period (−495 to −165 ms before the press, SVM 1b) in matched-latency trials, indicating that this subpopulation did not reliably encode movement type prior to lever press onset.
(J) Same as (E), for cluster 3 neurons
(K) Same as (F), for cluster 3 neurons
(L) Same as (G), for cluster 3 neurons

Shaded area in all line plots is ± SEM

**Figure S5**

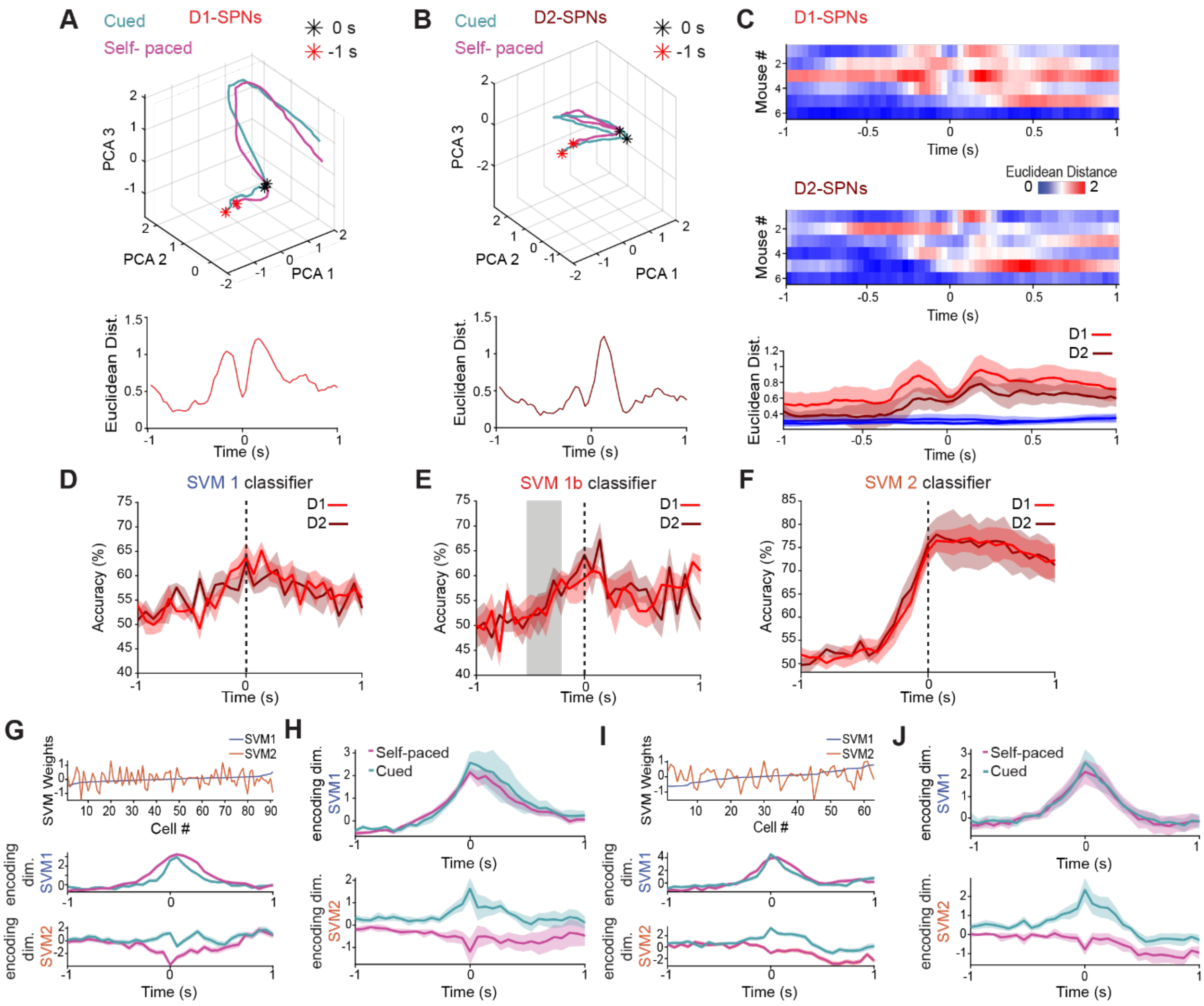


**Figure S5**, related to Figure 4: D1- and D2-SPNs exhibit similar action-encoding population dynamics

(A) Top: Principal component analysis of D1-SPN population activity shows that neural trajectories for cue-evoked (teal) and self-paced (magenta) trials in an example mouse. Black asterisk is time of the press/reward delivery and red asterisk is 1 second before that. Bottom: distance between neural trajectories shown above versus time.

(B) Same as (A), for D2-SPNs

(C) Euclidean distance between neural trajectories for cue-evoked and self-paced trials is shown for individual mice in D1-SPN (top) and D2-SPN (middle) populations. Bottom: The average of all D1-SPNs in all mice (red) and D2-SPNs (dark red) in all mice compared to shuffled data (blue).

(D) Support vector machine (SVM) classifier trained on 90% of the data and tested on the remaining 10% successfully predicted whether lever presses were cue-evoked or self-initiated based on activity in both D1-SPNs (red) and D2-SPNs (dark red) populations, indicating that both cell types carry distinguishable population activity patterns across trial types.

(E) SVM classification (SVM 1b) accuracy rose above chance during the cue period (−495 to −165 ms before the press) for matched-latency trials for both D1-SPNs (red) and D2-SPNs (dark red).

(F) For D1-SPNs (red) and D2-SPNs (dark red) neurons, the classifier distinguishing lever presses from baseline (SVM2) performed above chance before movement onset, indicating that this ensemble contributes to a shared preparatory neural dimension across both action types.

(G) For D1-SPNs, top: weights from SVM2 (press vs. baseline) are plotted in order of magnitude (red), with overlaid weights from SVM1 (cue vs. self-paced) showing distinct and uncorrelated contributions to each classification axis. Projections of D1 SPN activity onto these SVM-defined axes reveal trial-type-specific separation peaking at movement onset (middle) and shared motor-related dynamics across conditions (bottom).

(H) Same as G middle and bottom, for all D1-SPNs

(I) Same as G, for D2-SPNs

(J) Same as (I) middle and bottom, for all D2-SPNs

Shaded bars in all line plots are ± SEM.
